# Machine learning predicts rapid relapse of triple negative breast cancer

**DOI:** 10.1101/613604

**Authors:** Yiqing Zhang, William Nock, Meghan Wyse, Zachary Weber, Elizabeth Adams, Sarah Asad, Sinclair Stockard, David Tallman, Eric P. Winer, Nancy U. Lin, Mathew Cherian, Maryam B. Lustberg, Bhuvaneswari Ramaswamy, Sagar Sardesai, Jeffrey VanDeusen, Nicole Williams, Robert Wesolowski, Daniel G. Stover

## Abstract

**Purpose:** Metastatic relapse of triple-negative breast cancer (TNBC) within 2 years of diagnosis is associated with particularly aggressive disease and a distinct clinical course relative to TNBCs that relapse beyond 2 years. We hypothesized that rapid relapse TNBCs (rrTNBC; metastatic relapse or death <2 years) reflect unique genomic features relative to late relapse (lrTNBC; >2 years).

**Patients and Methods:** We identified 453 primary TNBCs from three publicly-available datasets and characterized each as rrTNBc, lrTNBC, or ‘no relapse’ (nrTNBC: no relapse/death with at least 5 years follow-up). We compiled primary tumor clinical and multi-omic data, including transcriptome (n=453), copy number alterations (CNAs; n=317), and mutations in 171 cancer-related genes (n=317), then calculated published gene expression and immune signatures.

**Results:** Patients with rrTNBC were higher stage at diagnosis (Chi-square p<0.0001) while lrTNBC were more likely to be non-basal PAM50 subtype (Chi-square p=0.03). Among 125 expression signatures, five immune signatures were significantly higher in nrTNBCs while lrTNBC were enriched for eight estrogen/luminal signatures (all FDR p<0.05). There was no significant difference in tumor mutation burden or percent genome altered across the groups. Among mutations, only *TP53* mutations were significantly more frequent in rrTNBC compared to lrTNBC (Fisher exact FDR p=0.009). To develop an optimal classifier, we used 77 significant clinical and ‘omic features to evaluate six modeling approaches encompassing simple, machine learning, and artificial neural network (ANN). Support vector machine outperformed other models with average receiver-operator characteristic area under curve >0.75.

**Conclusions:** We provide a new approach to define TNBCs based on timing of relapse. We identify distinct clinical and genomic features that can be incorporated into machine learning models to predict rapid relapse of TNBC.

## INTRODUCTION

Triple negative breast cancer (TNBC) is an aggressive breast cancer subtype defined by lack of targetable estrogen receptor (ER), progesterone receptor (PR) and HER2.^1^ TNBC accounts for 15% of breast cancer cases yet is responsible for 35% of breast cancer related deaths.^1,2^ TNBCs tend to present with higher grade, large size, and often involve lymph nodes at diagnosis. ^3^ TNBCs are more likely to develop distant rather than local recurrence compared to hormone receptor-positive counterparts and spreads more frequently to lung and brain and less frequently to bone.^2,4^ Understanding determinants of distant relapse is imperative as the median overall survival after diagnosis of metastatic disease was historically only 13-17 months^2,5^ and even among patients with PD-L1 positive TNBC receiving chemo-immunotherapy remains 25 months.^6^

The existing TNBC subsets/groupings provide a critical framework for understanding <u>intrinsic genomic characteristics</u> but are only associated with modest differences in patient survival. Clinically, we know that the majority of patients diagnosed with TNBC are long-term survivors.^1,2,7^ Among the 20-30% of TNBCs who develop metastatic disease, a subset have an aggressive phenotype associated with rapid relapse, therapeutic resistance, and poor prognosis while others have a relatively late relapse, associated with more indolent or treatment responsive disease.^7^ To more accurately understand the differences in patient outcome in TNBC, we sought to investigate TNBCs defined by timing of outcome: rapid versus late versus no relapse.

Advances in sequencing technology have facilitated comprehensive molecular profiling of breast cancers, including subsets such as TNBC.^8,9^ A major challenge for TNBC is inter-tumor heterogeneity, thought to be an important contributor to therapeutic resistance. A landmark transcriptional analysis of over 500 primary TNBCs revealed six subtypes of TNBC with distinct expression profiles.^10,11^ An integrated genomic analysis of TNBCs presented a unifying theory with four main TNBC subsets: 1) basal-like, immune-activated; 2) basal-like, immune suppressed; 3) luminal androgen receptor subtype: non-basal subtype with few CNAs; 4) mesenchymal: genomically unstable with fibroblast/EMT phenotype.^12^ In addition, genomic analyses demonstrate high frequency of *TP53* mutation, present in more than 75% of TNBCs, as well as mutations in *PIK3CA* in approximately 25% of cases.^12–14^ TNBCs also reflect widespread copy number alterations, suggesting that lack of genomic integrity is a key mutational process in TNBC.^12–14^

Clinical datasets may be limited in size, so accurate predictive modeling is crucial in a disease such as TNBC that has multiple distinct subsets.^15^ To be successful, predictive model needs to be able to learn from a small input and extract high-level, accurate information.^15^ There is growing implementation of various machine learning approaches to many areas of research including molecular subtypes of breast cancer^16,17^ and pathologic image analyses, such as detection of lymph node metastases in women with breast cancer.^18^ Determining the best approach for predictive modeling depends on size of data set, type of features, and outcome.^19^

In this study, we identify distinct genomic features among primary TNBCs categorized based on outcome: rapid (rrTNBC), late (lrTNBC) and no relapse (nrTNBC). Using a comparative modeling approach, we show that machine learning approaches provides an optimal predictor of rapid relapse for primary TNBCs.

## METHODS

### Patient and Tumor Characteristics

Patient-specific data were obtained from TCGA^13^, METABRIC^20,21^, and from our published meta-analysis (as described previously).^8^ These variables included age at diagnosis, grade, stage at diagnosis, pathologic receptor status (estrogen receptor (ER), progesterone receptor (PR), and HER2), response to neoadjuvant chemotherapy (where available), and distant metastasis-free or overall survival. Triple-negative breast cancer was defined as being negative for ER, PR, and HER2.Pathologic receptor status was ‘positive’ or ‘negative’ based on the following definitions: for ER and PR immunohistochemistry (IHC), 0 was defined as ‘negative’, 2-3 was defined as ‘positive’, and an IHC of 1 was considered indeterminate. For HER2, a FISH HER2/CEP17 ratio of greater than 2.0 was defined as ‘positive’. Chemotherapy response was categorized as pathologic complete response (pCR) or residual disease (RD) based on study-reported outcomes.

### Genomic Data

For data from the Molecular Taxonomy of Breast Cancer International Consortium (METABRIC), normalized gene expression data, copy number data, and somatic mutation data for 171 cancer-related genes were obtained from the publicly available European Genome-phenome Archive (IDs EGAD00010000210 and EGAD0001000021) and associated publications^21,22^. Copy number segmented data files were processed using GISTIC2.0^23^. For data from TCGA, breast cancer gene expression data, GISTIC copy number data, and somatic mutation data were obtained from the UCSC cancer browser (now XENAbrowser; version 2015-02-24). Gene expression data from published studies of breast cancer patients prior to neoadjuvant chemotherapy were compiled as previously described.^8^

### Gene Expression Signatures, Expression-Based Subtypes, and Inferred Immune Subsets

Given gene expression data from multiple studies and disparate platforms, gene expression data for all TNBCs for each dataset (METABRIC n=287, TCGA n=160, neoadjuvant dataset n=446) were extracted and quantile normalized within TNBCs from each study then median centered. Due to complexities of comparing single gene expression or differential expression across datasets/platforms/batches, we only evaluated summary expression metrics (e.g. signatures, intrinsic subtypes, CIBERSORT proportions). Gene expression signatures were compiled from published studies, as previously described.^8^ We determined PAM50 intrinsic breast cancer subtype using the ‘Bioclassifier’ package from Parker et al after balancing TNBC data with an equal number of ER-positive cases for each dataset.^24^ Triple-negative breast cancer (TNBC) subtype was determined using the TNBCtype tool.^10,25^ Proportion of infiltrating immune cell subsets were calculated using the CIBERSORT algorithm.^26^

### Modeling

We compared the performance of a multinomial logistic regression model to Artificial Neural Network (ANN), K-Nearest Neighbors (KNN), Support Vector Machine (SVM), Linear Determinant Analysis (LDA), and Random Forest (RF). Multinomial logistic regression model is an extension of logistic regression model that allows for a dependent variable with more than two levels. ANNs are based on a collection of ‘neurons’ or ‘nodes’ arranged in layers and features are fed from one layer to the next with each layer extracting different high-level features. The last hidden layer forms a high-level representation to make a classification decision. The KNN algorithm classifies data points based on other most similar data points (‘nearest neighbors’). SVM develops an optimal hyper-plane in the feature space to separate different outcomes. LDA compares the log of the estimated density function, called the discriminant function, and the class assigned is the one with the highest value of the discriminant function. RF develops many random decision trees and assigns each new data point based on majority votes among all decision trees.

To adjust the magnitude of features for optimization algorithms (often used in SVM and ANN),^27^ all numerical attributes were normalized. We identified 148 features significantly different between rrTNBC, lrTNBC, and nrTNBC (nominal p < .05) then used the findCorrelation function in caret R package^28^ to remove highly correlated numerical features (correlation > .85) or highly associated features (Cramer’s V > .5) to avoid high dimensionality, resulting in a final model input using 77 features. To have adequate data to train our models and sufficient testing samples to evaluate model performance, we partitioned our sample into 70% training set and 30% test set. 10-fold validation was performed on the 70% training data using caret^28^ and h2o R packages,^29^ and models were tested on the 30% test data.

We tuned each model to optimize performance. We used lasso reduction for multinomial logistic regression model, and tuned the regularization parameter lambda. The chosen set of lambda is an equally spaced array of 20 values from 1e-5 to .1. The ANN model was tuned on number of hidden layers, neurons on each hidden layer, and activation function using h2o.deeplearning function. The structure we chose were 1 to 3 hidden layers consisting of 5, 10, or 20 neurons on each layer with either Rectifier or Tanh activation function leading to the final output. We tuned number of nearest neighbors of the KNN model from 1 to 15 while trying to avoid multiples of 3. Built-in tuning was used for the LDA, SVM, and RF models. The final optimal model was chosen based on the logloss metric (-log(likelihood function)).

### Model Performance

To evaluate model performance, we assessed each group as a positive outcome (rrTNBC versus all others; lrTNBC vs all others; nrTNBC vs all others) using receiver-operator characteristic (ROC) curves. We used multi_roc function in multiROC^30^ to compute micro-average area under the curve (AUC) of three ROC curves to evaluate the overall performance of the models. To avoid sampling bias, we calculated the average AUC on the 30% test data for 10 independent runs.

### Statistical Analysis

All statistical analyses were performed in R version 3.4.1. Contrasts in patient and tumor characteristics were evaluated using Pearson chi-squared tests. The association of signatures to neoadjuvant chemotherapy response was evaluated using simple linear regression and t-tests. All calculations of association were multiple-testing corrected using the Benjamini–Hochberg procedure for false discovery rate. For continuous variables, we calculated p-values comparing rapid vs. late and relapse vs. no relapse using ANOVA and logistic regression. For count variables (e.g. mutated versus not) we used Fisher exact test to evaluate relapse vs. not and rapid vs. late relapse. P-values for CIBERSORT and mutation signatures were evaluated using logistic regression, and for CNAs and mutations were evaluated using Fisher exact tests. Data visualization was made using ggplot2.^31^

## RESULTS

### Defining Rapid versus Late versus No Relapse Triple-Negative Breast Cancer

From three large cohorts with primary breast cancer genomic data – TCGA,^13^ METABRIC,^20,21^ and our prior breast cancer gene expression meta-analysis^8^ - we identified 893 TNBCs from a total of 4473 breast cancer cases. For our analyses, we included patients with at least 60 months of follow-up or those with a distant metastasis-free survival (DMFS) event prior to our 60-month cutoff, leaving a total of 453 TNBCs in our evaluable dataset. Of these, 453 had gene expression data, 317 had copy number data, and 317 had mutation data. (**Figure 1A**).

**Figure 1.**
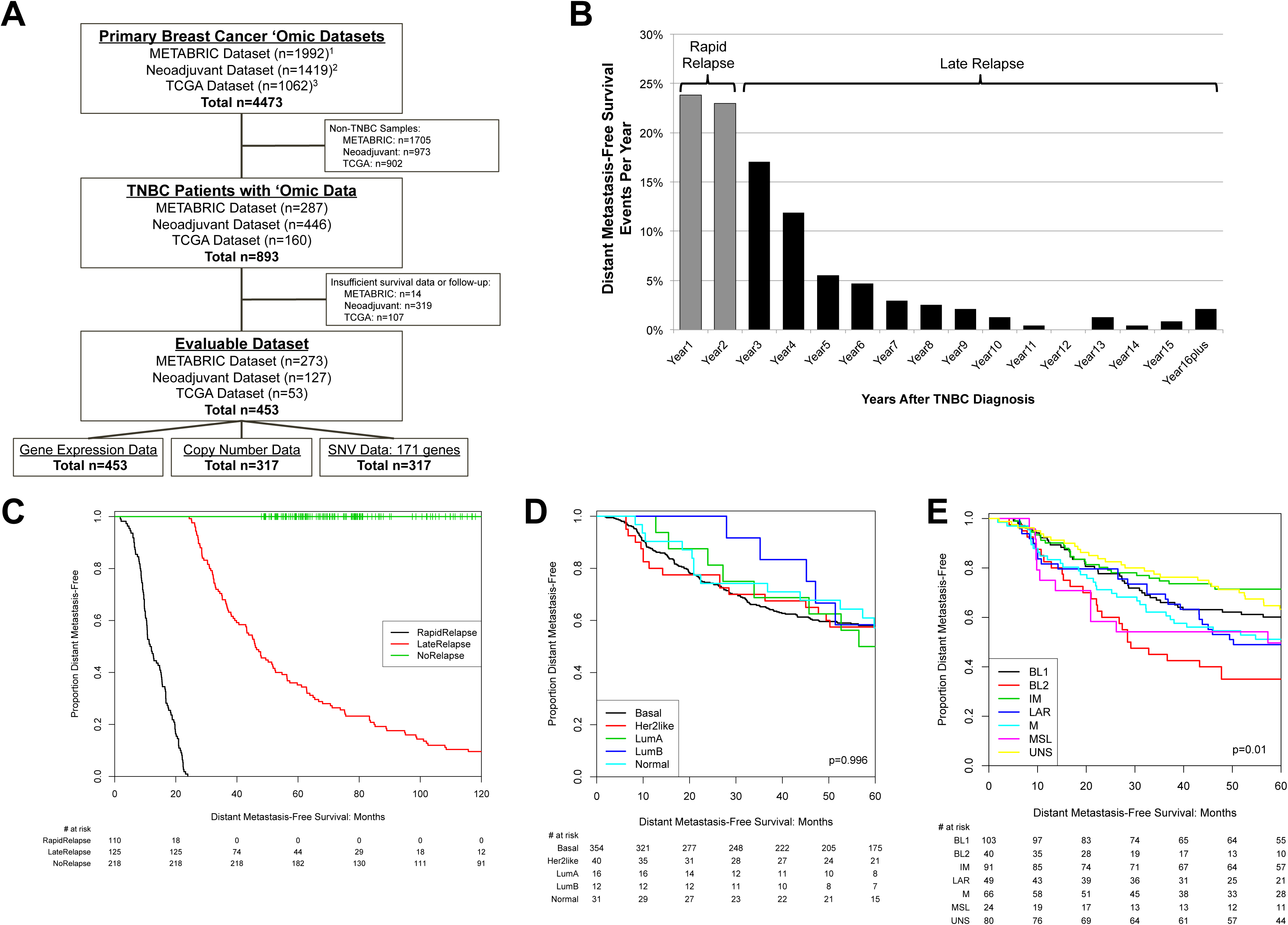
Study design and definition of triple-negative breast cancer (TNBC) rapid versus late relapse. **(A)** REMARK diagram. **(B)**. Proportion of distant metastasis-free survival (DMFS) events per year after diagnosis among evaluable dataset. ‘Rapid relapse’ was defined as DMFS events within the 2 years of diagnosis and ‘late relapse’ DMFS events beyond 2 years. **(C-E)** Kaplan-Meier diagram of DMFS in study cohort reflecting TNBC group definitions **(C)**, compared with DMFS by intrinsic subtype approaches PAM50 subtype **(D)**, and Lehmann TNBC subtype **(E).** P-value indicates log-rank test.

We assessed the percentage of total DMFS events each year (**Figure 1B**). In this dataset, over 20% of DMFS events occurred within the first two years after diagnosis, categorized as ‘rapid relapse’ (rrTNBC). While most DMFS events occurred within the first five years after diagnosis, 1-3% of events occurred annually in years 6-10 with sporadic events beyond 10 years (lrTNBC). Our main goal was to identify differences among TNBCs with clinically distinct outcomes, so we visualized DMFS for our relapse categorization (**Figure 1C**) in comparison with DMFS for existing intrinsic expression-based subtype approaches PAM50^24^ (**Figure 1D**) or Lehmann/Pietenpol TNBCtype^10^ (**Figure 1D**) within the same cohort. The Lehmann/Pietenpol TNBCtype (log-rank p=0.01) but not PAM50 was associated with significant differences in DMFS. The strikingly different visualized outcomes suggested that our relapse categorization does, in fact, identify truly distinct subsets based on outcome when compared to approaches that focus on intrinsic features.

### Patient and Tumor Characteristics

We evaluated the association of clinical, pathologic, and intrinsic expression subtype with rapid vs. late vs. no relapse status (**Table 1**). There was no significant difference in age at diagnosis or grade; however, rrTNBCs were significantly more likely to be higher stage (Chi-square p=1.9e-10). The majority of patients were basal-like PAM50 subtype (78%); however, lrTNBCs were significantly more likely to be non-basal (non-basal: rrTNBC 18%, lrTNBC 29%, nrTNBC 20%, Chi-square p=0.03). Lehmann/Pietenpol TNBC subtype also reflected significant differences across groups (Chi-square p=0.02), although each subtype was represented within each relapse category: the immunomodulatory phenotype was highest in nrTNBC (16% rrTNBC, 16% lrTNBC, 24% nrTNBC),luminal androgen receptor was highest in lrTNBC (9% rrTNBC, 16% lrTNBC, 9% nrTNBC), and basal-like 2 was highest in rrTNBC (15% rrTNBC, 9% lrTNBC, 6% nrTNBC). A subset of patients in this cohort (127/453; 28.0%) had data on response to neoadjuvant chemotherapy (NAC). As anticipated, those patients with rrTNBC or lrTNBC were significantly more likely to have residual disease (RD) after neoadjuvuant chemotherapy (93% and 94% RD, respectively), relative to those with nrTNBC (51% RD; Chi-square p=1.9e-7).

**Table 1.**
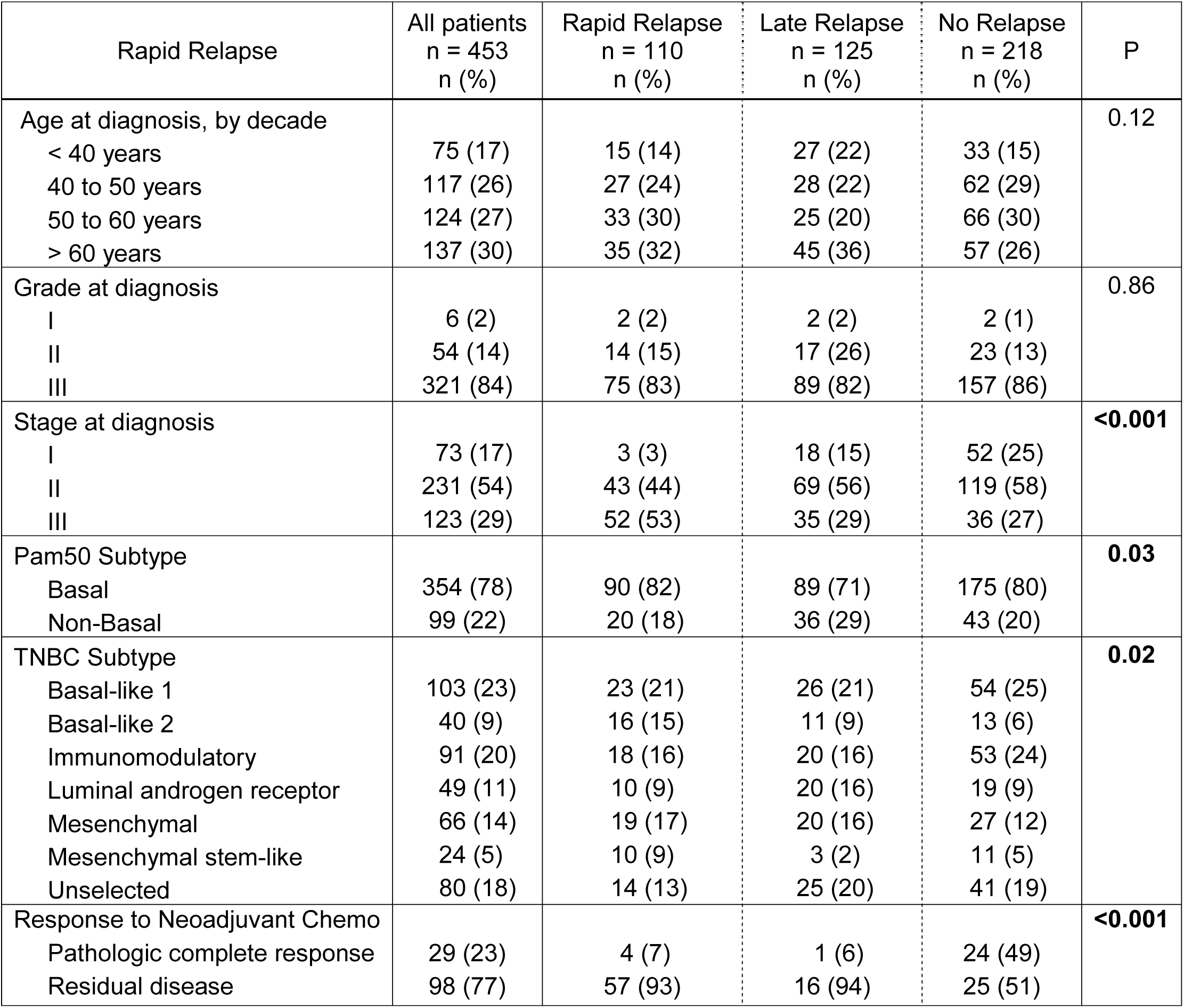
Cohort clinical and pathologic features

### Response to Neoadjuvant Chemotherapy and Survival in TNBC: Immune and Expression Signatures

Response to NAC is known to be a robust prognostic biomarker in TNBC^32^ and, in this cohort, RD after NAC was among the clinicopathologic features most strongly associated with rapid and late versus no relapse (**Table 1**). The patients with data on response to NAC all had whole transcriptome data but no available mutation or copy number data, so we sought to understand expression features associated with NAC response and DMFS. We calculated a score for 125 published gene expression signatures and evaluated the association of each signature with NAC response (pathologic complete response versus RD) by simple linear regression and hazard ratio for each signature using DMFS. Signatures were grouped by phenotype as previously described^8^ (n=127 patients; **Figure 2A**). Immune signatures were associated with better prognosis and most were also associated with improved response to NAC. Proliferation signatures tended to be associated with improve*ed response to NAC, as we have previously described^8^, yet there was variable association with DMFS. Estrogen receptor/HER2 signatures were associated with poor response to NAC but better DMFS. Mesenchymal signatures tended to be associated with worse DMFS without clear association with response to NAC.

**Figure 2.**
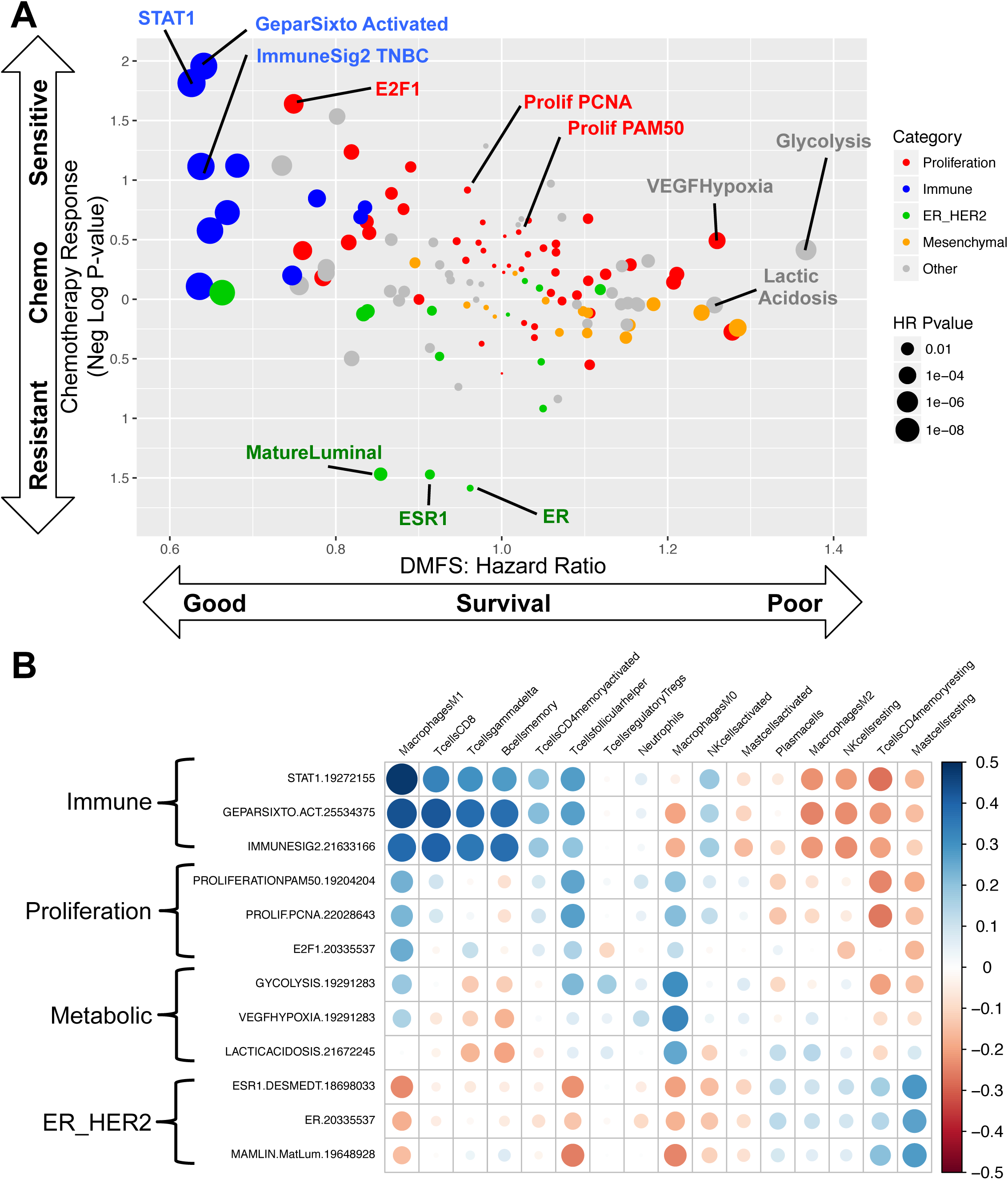
Immune and Expression Signatures and Response to Neoadjuvant Chemotherapy and Survival in TNBC. **(A)** The calculated score for 125 published gene expression signatures for 127 patients with data on response to neoadjuvant chemothrapy and distant metastasis-free survival (DMFS). Each signature is a point. The association of each signature with neoadjuvant chemotherapy response (pathologic complete response versus RD) by simple linear regression (y-axis) and hazard ratio for each signature using DMFS (x-axis) are displayed. Signatures were grouped by phenotype (as previously described^8^), identified by color: proliferation signatures (red), immune signatures (blue), ER/HER2 signatures (green), mesenchymal signatures (orange), others (grey). Size of each point relates to the hazard ratio p-value for each signature. **(B)** The association of three representative signatures from each group (immune, proliferation, ER/HER2, mesenchymal) with the relative proportion of 22 inferred immune cell subsets via CIBERSORT across all samples with gene expression data (n=453) are visualized using CorrPlot.^26,62^

To understand what immune cell types in the tumor microenvironment may be reflected by the immune signatures, we evaluated the association of three representative signatures from each group (immune, proliferation, ER/HER2, mesenchymal) with the relative proportion of 22 inferred immune cell subsets via CIBERSORT (**Figure 2B**).^26^ Immune signatures were strongly positively correlated with anti-tumor immune cell types including M1 macrophages, CD8 T-cells, and memory B-cells (all Pearson’s r >= 0.3, all p<1.2e-8) and anti-correlated with immune suppressive cell types including M2 macrophages, memory resting CD4 T-cells, resting NK cells, and resting mast cells. ER/HER2 signatures reflected an almost opposite pattern to immune signatures, with positive correlation to immune suppressive cell types and anti-correlation with anti-tumor immune cell type. Proliferation signatures had a similar pattern to immune signatures, although less striking, while metabolic signatures appeared to have a strong correlation specifically with M0 macrophages (all Pearson’s r>0.27, all p<8.4e-9). As a sensitivity analysis, we evaluated the association of three representative signatures from each group with 7 immune cell-type specific signatures from MSigDB^33,34^ (instead of CIBERSORT) and found similar results (Supplementary Figure 1A).

### Expression Signatures in Rapid versus Late versus No Relapse TNBC

To assess pathways and phenotypes associated with rapid vs. late vs. no relapse, a score was calculated for 125 published gene expression signatures across the entire dataset. Visual observation of all signatures or only the quarter with the greatest variance did not yield any clear patterns (Supplementary Figure 1B). Evaluating each signature individually across the three groups revealed 16 signatures that were significantly different (ANOVA FDR p<0.05; **Figure 3**, Supplementary Figure 2A-B). Among these, five signatures were immune-related: two from the GeparSixto trial^35^, an immune signature from TNBC subtypes,^10^ and two STAT1 signatures.^36,37^ All were significantly higher in nrTNBC than rrTNBC and lrTNBC. Eight of the sixteen significant signatures were related to estrogen receptor signaling and/or luminal phenotype – all were highest in lrTNBC, lowest in rrTNBC, and intermediate in nrTNBC. Finally, three signatures did not fall into a specific category and revealed mixed patterns of association. Most CIBERSORT immune subsets were not statistically significant (Supplementary Figure 2C), however, neutrophils were significantly higher in rrTNBC (ANOVA FDR p=0.001) while resting mast cells were significantly lower in lrTNBC (ANOVA FDR p=0.003).

**Figure 3.**
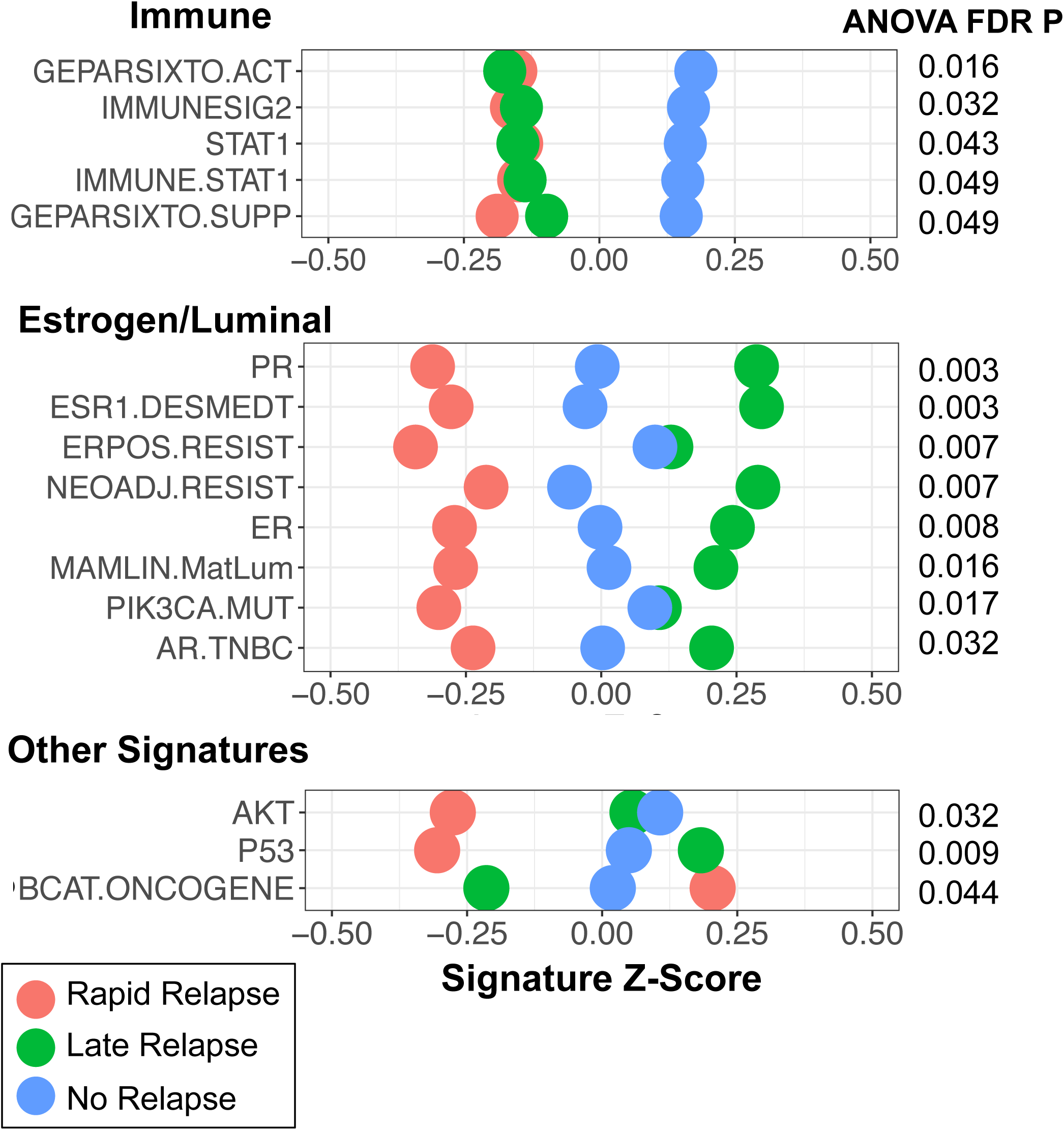
Expression Signatures in Rapid versus Late versus No Relapse TNBC. The calculated score for 16 published gene expression signatures that demonstrated statistical significance (ANOVA FDR p<0.05) comparing rapid versus late versus no relapse. Signatures visualized as relative values (Z-score) with rapid relapse (red), late relapse (green), and no relapse (blue).

### Mutations and Copy Number Alterations

In this cohort, 70% (317/453) of patients had data on single nucleotide variant/mutation data on 171 cancer-related genes and whole genome CNAs.^21^ Only a small subset of patients (11.7%; 53/453) had whole exome mutation data so we focused on the 171 cancer-related genes to ensure adequate statistical power. When evaluating general mutational features, there was no significant difference in mutations per megabase (ANOVA p=0.64; **Figure 4A**) nor percent genome altered by copy number (ANOVA p=0.96; **Figure 4B**).

**Figure 4.**
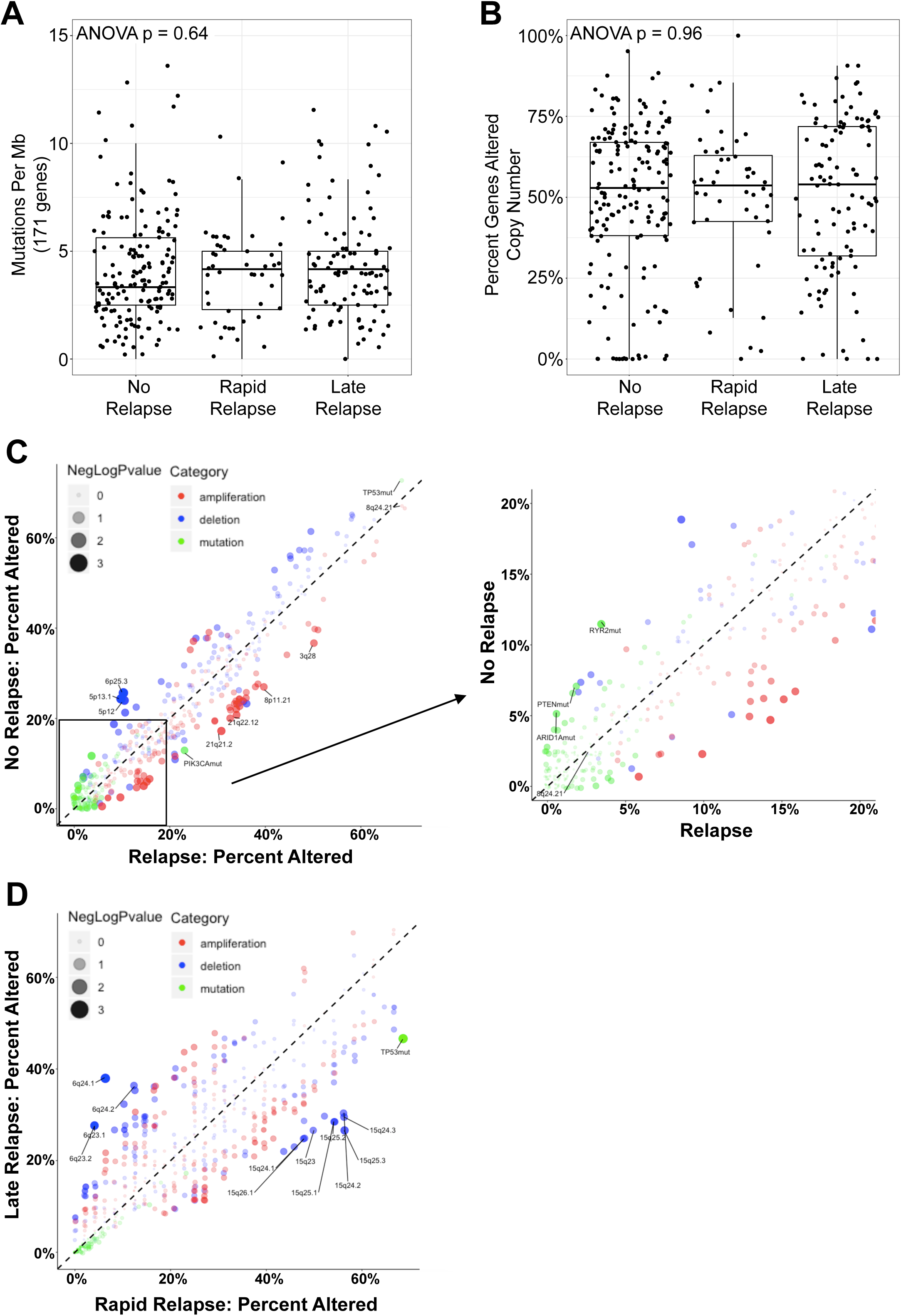
Mutations and copy number alterations in rapid versus late versus no relapse TNBCs. **(A)** Mutations per megabase of 171 cancer-related genes. **(B)** Percent genes altered by copy number gain (GISTIC 1 or 2) or loss (GISTIC -1 or -2). **(C)** Frequency of alteration of 171 cancer-related genes (green dots), copy number gains (red dots) or losses (red dots) by cytoband among rapid relapse (x-axis) versus no relapse (y-axis) TNBCs (**C)** or rapid relapse (x-axis) versus late relapse (y-axis) TNBCS **(D)**. Size of dot indicates negative log of p-value for Fisher exact test with those genes and cytobands indicated demonstrate nominal p<0.05. Zoomed-in image of those alterations with <20% frequency indicated in right panel.

We first compared the frequency of alteration for each mutation and cytoband (for CNAs) for relapse (rrTNBC + lrTNBC) versus nrTNBC (**Figure 4C**) given low frequency of mutation for most genes. There were no genes that were significantly different after multiple testing when comparing relapse versus no relapse but *PIK3CA* mutations were more frequent in relapse relative to nrTNBC; and *PTEN, ARID1A,* and *RYR2* mutations enriched in nrTNBC relative to rrTNBC (Fisher exact nominal p<0.05). We then compared rrTNBC versus lrTNBC (**Figure 4D**) and found that rrTNBC were significantly more likely to harbor a mutation in *TP53* compared to lrTNBC patients (Fisher exact FDR p=0.009). There were no other genes that were significantly different when comparing rapid vs. late vs. no relapse after multiple test correction (Supplementary Figure 3A). Among CNAs, the copy number landscape was similar across all three groups (Supplementary Figure 3B) and there were no significantly altered genes or regions among rapid vs. late vs. no relapse after multiple test correction. There were several regions that demonstrated enrichment within specific groups (nominal p<0.05), including: 3q28 gain, 8p11 gain, and 21q21-22 gain enriched in relapse relative to nrTNBC; 5p12-13 loss and 6p25 loss enriched in nrTNBC relative to relapse; 6q23-24 loss enriched in lrTNBC relative to rrTNBC; and 15q23-26 loss enriched in rrTNBC relative to lrTNBC (**Figure 4C-D**).

### Optimal clinical and multi-‘omic model of rapid versus late versus no relapse in TNBC

Having identified discrete clinical, expression, immune, mutation, and copy number features among primary TNBCs with distinct clinical outcomes, we sought to develop an optimal, multi-‘omic predictive model for rrTNBC versus lrTNBC versus nrTNBC. As has been applied previously in complex breast cancer data^17^, we compared performance of multinomial logistic regression model to Artificial Neural Network (ANN), K-Nearest Neighbors (KNN), Support Vector Machine (SVM), Linear Determinant Analysis (LDA), and Random Forest (RF) (**Figure 5A**). We included any feature present in all samples (n=312) with a nominal p<0.05, which resulted in a total of 77 clinical, expression, immune, mutation, and copy number features (**Figure 5A**). Our total multi-‘omic dataset was divided into 70% training and 30% validation cohorts, independently sampled for each cross validation (10-fold cross validation). We assessed each model’s performance using three separate receiver-operator characteristics (ROCs) (rrTNBC as positive, lrTNBC as positive, and nrTNBC as positive) due to 3-level classification. We then integrated these into a single ROC curve via micro-averaging (**Figure 5B**). All models performed relatively well, with highest average AUC 0.772 (SVM) and lowest average AUC 0.681 (ANN) (**Figure 5C**). When comparing models, SVM had the highest average AUC, significantly higher than all other models (all Wilcoxon rank sum p<0.05), followed by RF and multinomial. As a sensitivity analysis, we repeated our approach using only the 18 most significant features that were all FDR p<0.05 in descriptive statistics and found that using fewer features led to similar average AUC (range 0.695 to 0.748), with random forest performing best (all Wilcoxon rank sum p<0.05) followed by SVM and multinomial (Supplementary Figure 3C).

**Figure 5.**
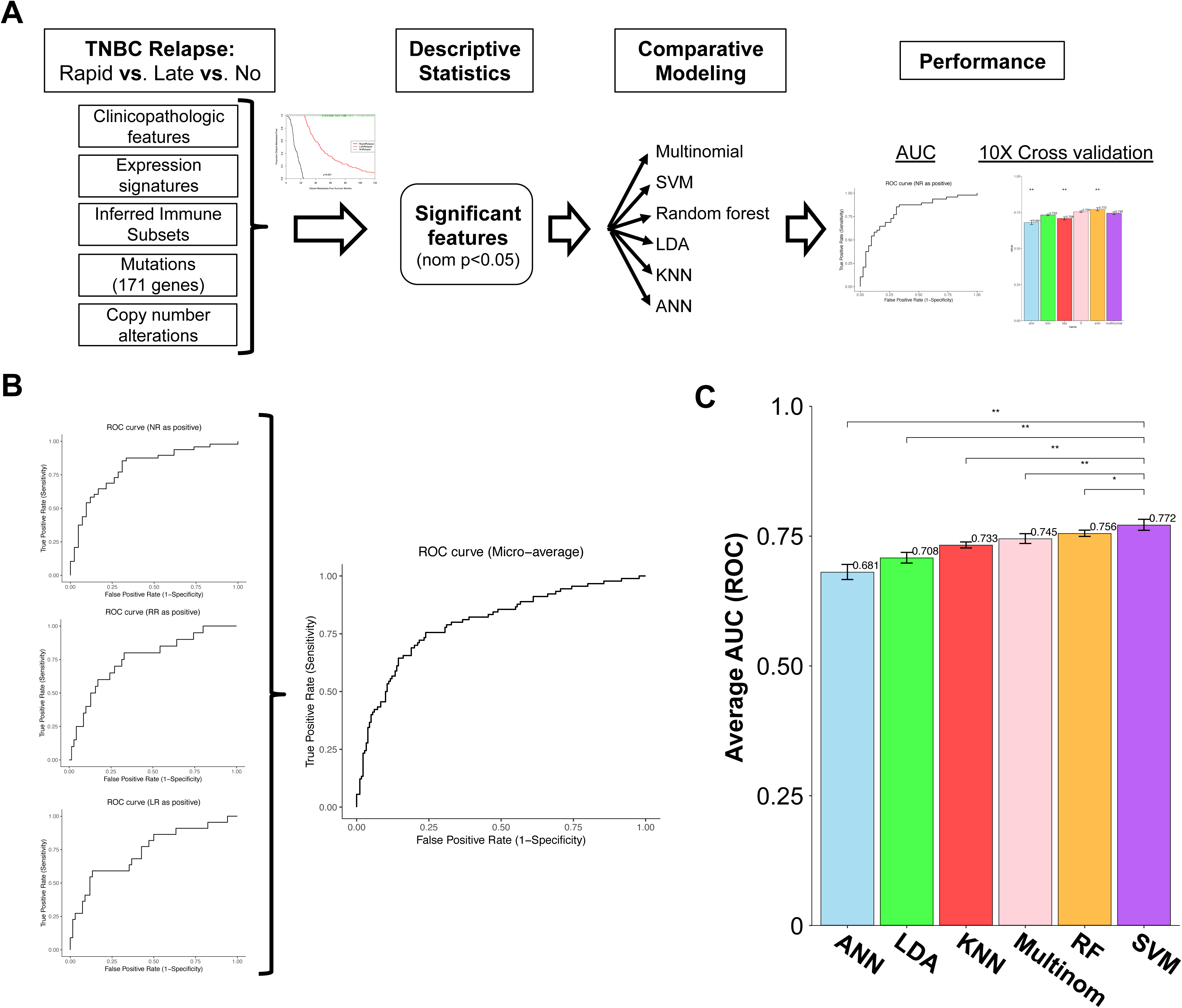
Developing an optimal clinical and multi-‘omic model of rapid versus late versus no relapse in TNBC. **(A)** Schematic of experimental steps including definition of variables, descriptive statistics, comparative modeling including model tuning, and assessment of model performance. **(B)** Example receiver-operator characteristic (ROC) plots for each model performance assessment using three separate ROCs (left panel; rrTNBC as positive, lrTNBC as positive, and nrTNBC as positive). These three ROCs were integrated these into a single ROC curve via micro-averaging (right panel) **(C)** Performance of Artificial Neural Network (ANN), Linear Determinant Analysis (LDA), K-Nearest Neighbors (KNN), multinomial logistic regression model (Multinom), Random Forest (RF), and Support Vector Machine (SVM). Our total dataset (n=312) was divided into 70% training and 30% validation cohorts, independently sampled for each cross validation (10-fold cross validation). Each model was tuned to ensure optimal performance within the constraints of a comparative modeling approach. Asterisks indicates significance by Wilcoxon rank sum, * indicates p<0.05 and ** indicates p<0.01.

## DISCUSSION

TNBC is a vexingly heterogeneous disease, as evidenced by multiple efforts to categorize subsets of which have primarily focused on intrinsic features of TNBC^10–12,24^ to reveal underlying biology, with fewer efforts focused on long-term outcomes. When considering clinical impact, response to neoadjuvant chemotherapy remains the best prognostic biomarker for TNBC,^32^ but once TNBC patients develop metastatic relapse, there are clear differences in disease course among TNBCs who develop relapse early versus late. To delineate these groups by relapse timing for further analyses, we provide a novel definition for rapid versus late relapse in TNBC and demonstrate clear differences in genomic features that cut across existing TNBC intrinsic subtypes.

When comparing TNBCs who relapse versus those who do not, both rrTNBC and lrTNBC had lower expression of immune signatures, reflecting reduced anti-tumor immune response compared with nrTNBCs. The correlation between immune cells and positive outcomes is well-established and tumor infiltrating lymphocytes (TILs) are associated with improved response to NAC as well as better prognosis in TNBC.^38–44^ Particularly given recent FDA approval of immunotherapy for metastatic TNBC^6^, there is great interest to augment the existing host anti-tumor immune response and also work to activate immune ‘cold’ tumors.^45–49^ The differences between late and rapid relapse primarily reflect differences in luminal features. lrTNBCs are more likely to be non-basal (primarily luminal A/B) and our data identify that eight estrogen/luminal signatures are associated with late relapse. Multiple groups have identified a ‘luminal androgen receptor’ subset of TNBC based on molecular classifications,^10,12^ and 40% (20/49) of the Lehmann LAR subtype tumors in our cohort ultimately had late relapse. The wide distribution of Lehmann TNBC subtypes among the lrTNBC group suggest that we may need additional approaches to identify tumors with this phenotype at risk for late relapse. Collectively, our data support the categorization by Burstein, et al^12^ and suggest that ‘rapid relpase’ TNBCs are likely enriched for the ‘basal-like immune suppressed’ phenotype.

Clinically, multi-‘omic models could serve as an adjunct to neoadjuvant chemotherapy response to predict patients at high risk of rapid relapse to guide more aggressive therapy while identifying those likely to be cured (even with residual disease) to avoid unnecessary toxicity of additional treatment. Given our interest to understand <u>what</u> genomic features contribute, we considered the best way to overcome the significant challenge extracting important features of machine learning algorithms. Because multi-‘omic data has clear features that can be extracted,^50^ we first identified the relatively few specific features that were significantly different across subsets (77 features from >35,000 initial data points) then built models based on *a priori* feature identification. This approach identified key genomic features up-front and led to overall good performance of multiple models but may miss unexpected interactions among raw features.

Stage at diagnosis was strongly associated with rapid relapse in univariate analyses and in the multinomial model (where feature contribution could be extracted), stage was the feature contributing most to the model. This suggests that despite our efforts to use complex biological data, stage remains critical. One hypothesis is that stage at diagnosis captures non-biological features such as socioeconomic or demographics, including income, education, insurance, and access to care.^51–55^ Race/ethnicity is complex,^56–58^ was largely unavailable in the included datasets, and warrants further study.^59,60^ We are currently evaluating the association of sociodemographic features and race/ethnicity with rrTNBC in other data sources.

While this study presents promising methods to categorize TNBC relapse it does possess some limitations. The retrospective nature of our data means that patients might not perfectly fit the definitions we assigned, for example, a small number of late relapse events may occur in the nrTNBC group after 5 years. We incorporated genomic data from multiple studies, generated using multiple platforms, and over years. While we have attempted to account for this through standard normalization approaches and analysis only of summary statistics (e.g. expression signatures not individual genes), batch/platform effects and computational analyses could impact our results. Therapy for TNBC has changed over the past 2-3 years, including widespread incorporation of capecitabine after neoadjuvant chemotherapy for residual disease based on CREATE-X^61^ and recent FDA approval of immunotherapy for metastatic, PD-L1 positive TNBC.^6^

In conclusion, we offer a new definition for ‘rapid relapse’ TNBC and provide evidence that rrTNBC reflects a distinct clinical entity characterized by unique genomic features. Predictive modeling may identify patients at high risk for ‘rapid relapse’ and offers potential to guide additional therapy or clinical trials for these high-risk patients.

## FIGURE LEGENDS

**Figure S1. Additional Analyses of Gene Expression Signatures. (A)** Sensitivity analyses of correlation between three representative signatures from each group (immune, proliferation, ER/HER2, mesenchymal) with the immune cell-specific signatures^33,34^ across all samples with gene expression data (n=453), visualized using CorrPlot.^26,62^ **(B)** Heatmap with hierarchical clustering of the gene expression signatures with the greatest variance (top 25%) across the dataset.

**Figure S2. Variation of Expression Signatures Across Rapid versus Late versus No Relapse Groups.** The calculated score for 16 published gene expression signatures that demonstrated statistical significance (ANOVA FDR p<0.05) comparing rapid versus late versus no relapse. The score value is presented for immune signatures **(A)** and estrogen/luminal signatures **(B)**. Each boxplot represents the 25^th^ to 75^th^ percentile with the median indicated as the central line and whiskers indicating 1.5 x interquartile range. **(C)** Immune cell subset proportion from CIBERSORT, visualized as relative values (Z-score) with rapid relapse (red), late relapse (green), and no relapse (blue).

**Figure S3. Mutation and Modeling Sensitivity Analyses**. **(A)** CoMut plot of gene-level mutation for the entire cohort, with mutation indicated in blue, visualized with ‘GenVisR’ package.^63^ **(B)** Frequency of gene-level copy number gains (red) or losses (blue) across the genome. **(C)** Sensitivity analysis of model performance for Artificial Neural Network (ANN), Linear Determinant Analysis (LDA), K-Nearest Neighbors (KNN), multinomial logistic regression model (Multinom), Random Forest (RF), and Support Vector Machine (SVM) using only 18 features (all FDR p<0.05). Asterisks indicates significance by Wilcoxon rank sum, ** indicates p<0.01.

