## Supplementary figures and images for "Machine learning predicts rapid relapse of triple negative breast cancer"

### Supplementary Figure 1

#### Supplementary Figure 1

**A**

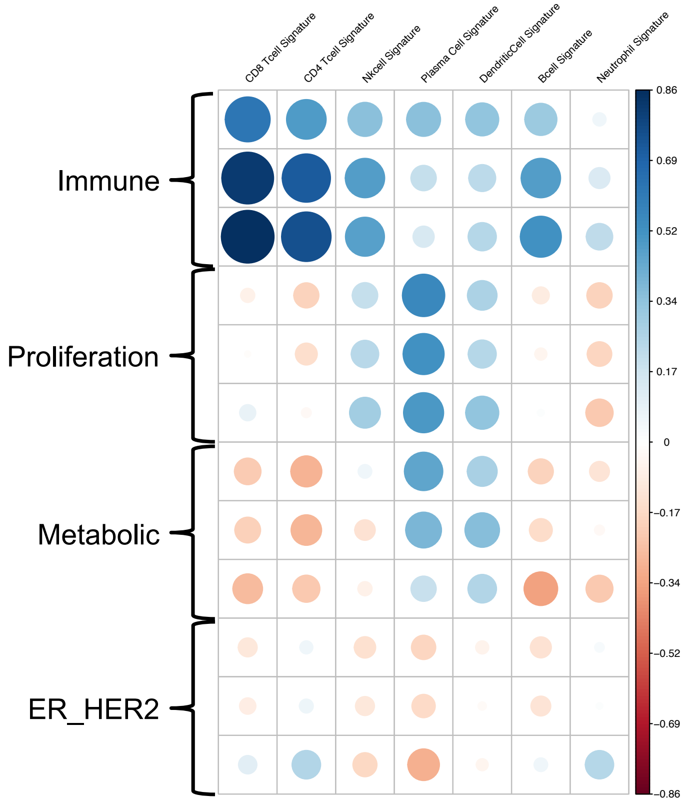


**B**

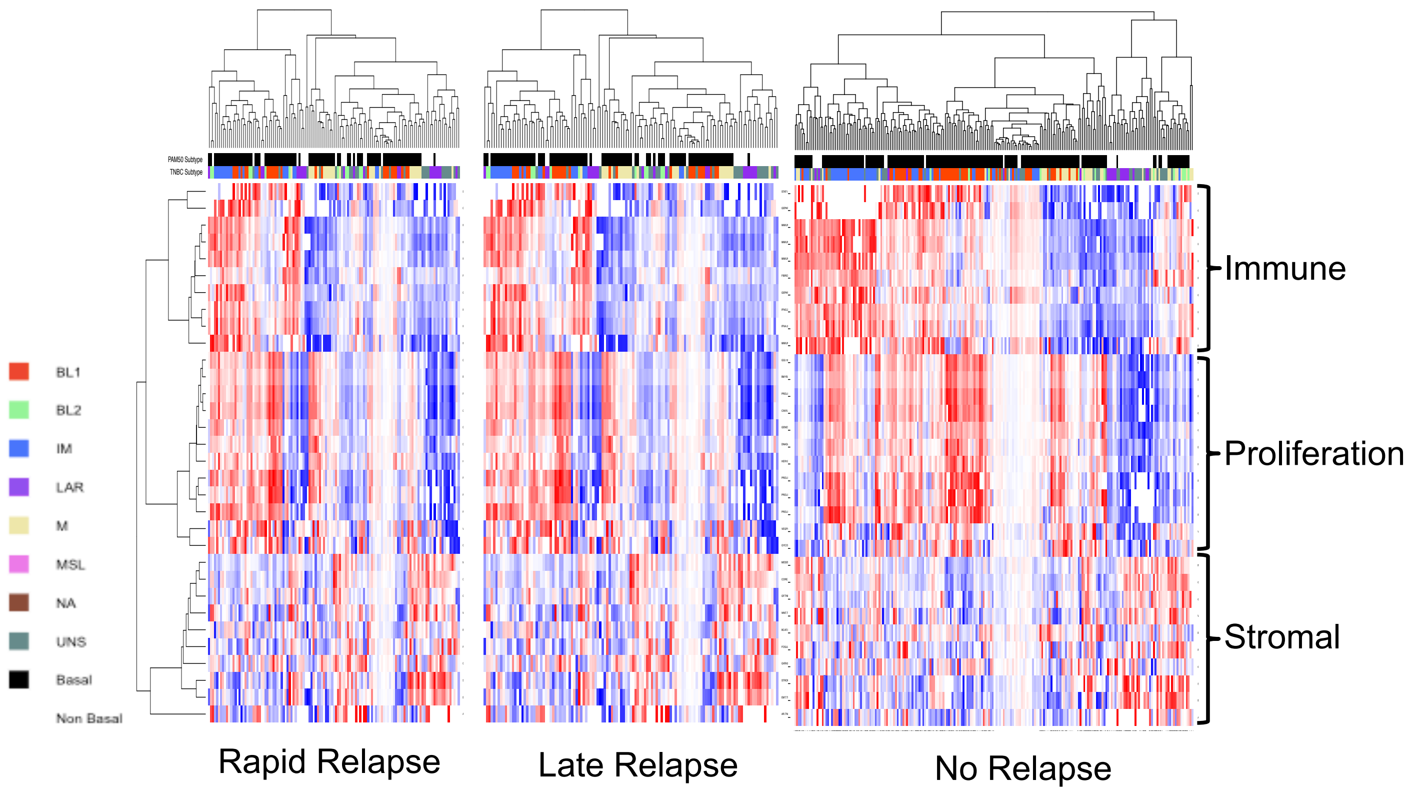

### Supplementary Figure 2

#### Supplementary Figure 2

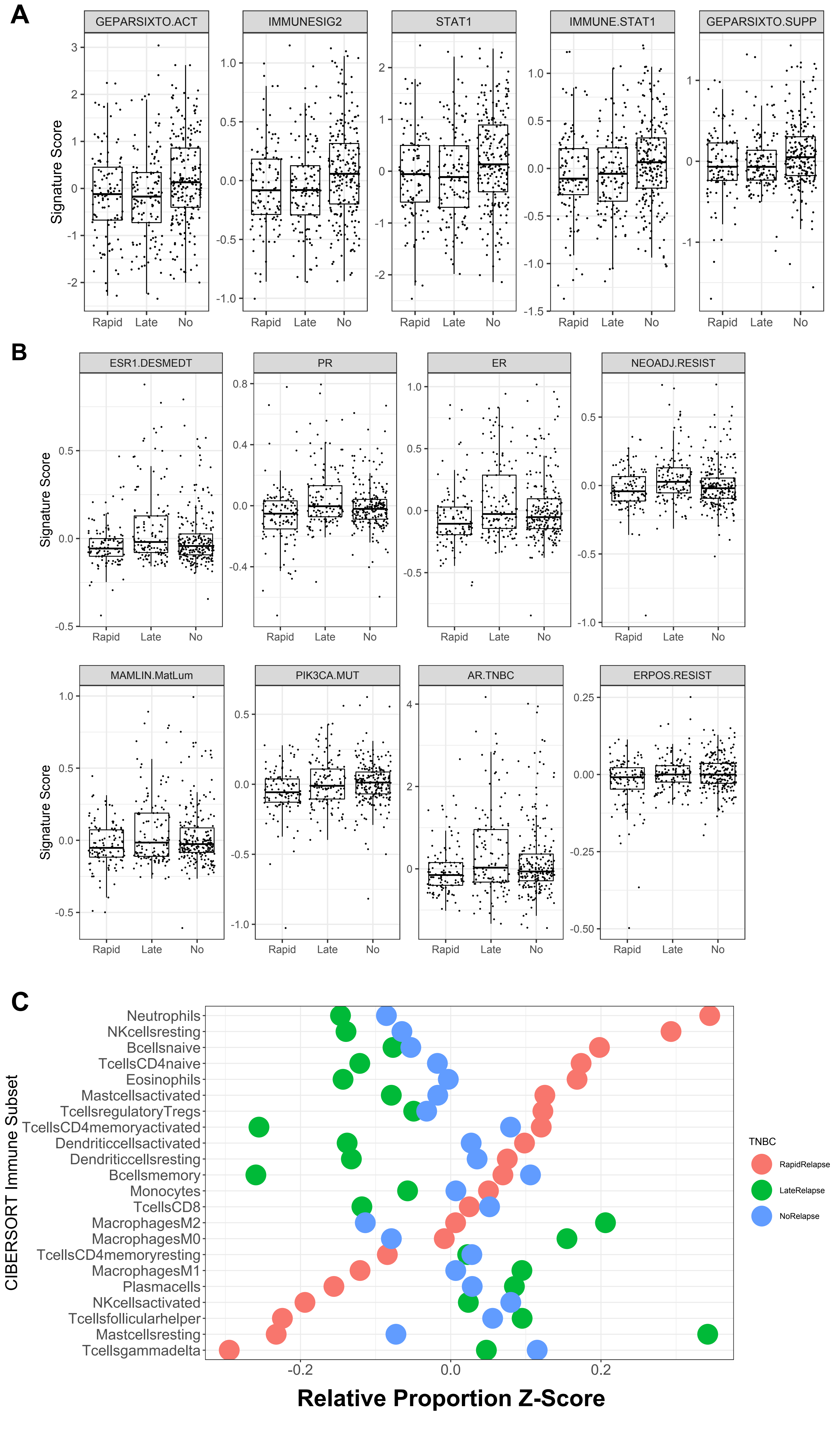

### Supplementary Figure 3

#### Supplementary Figure 3

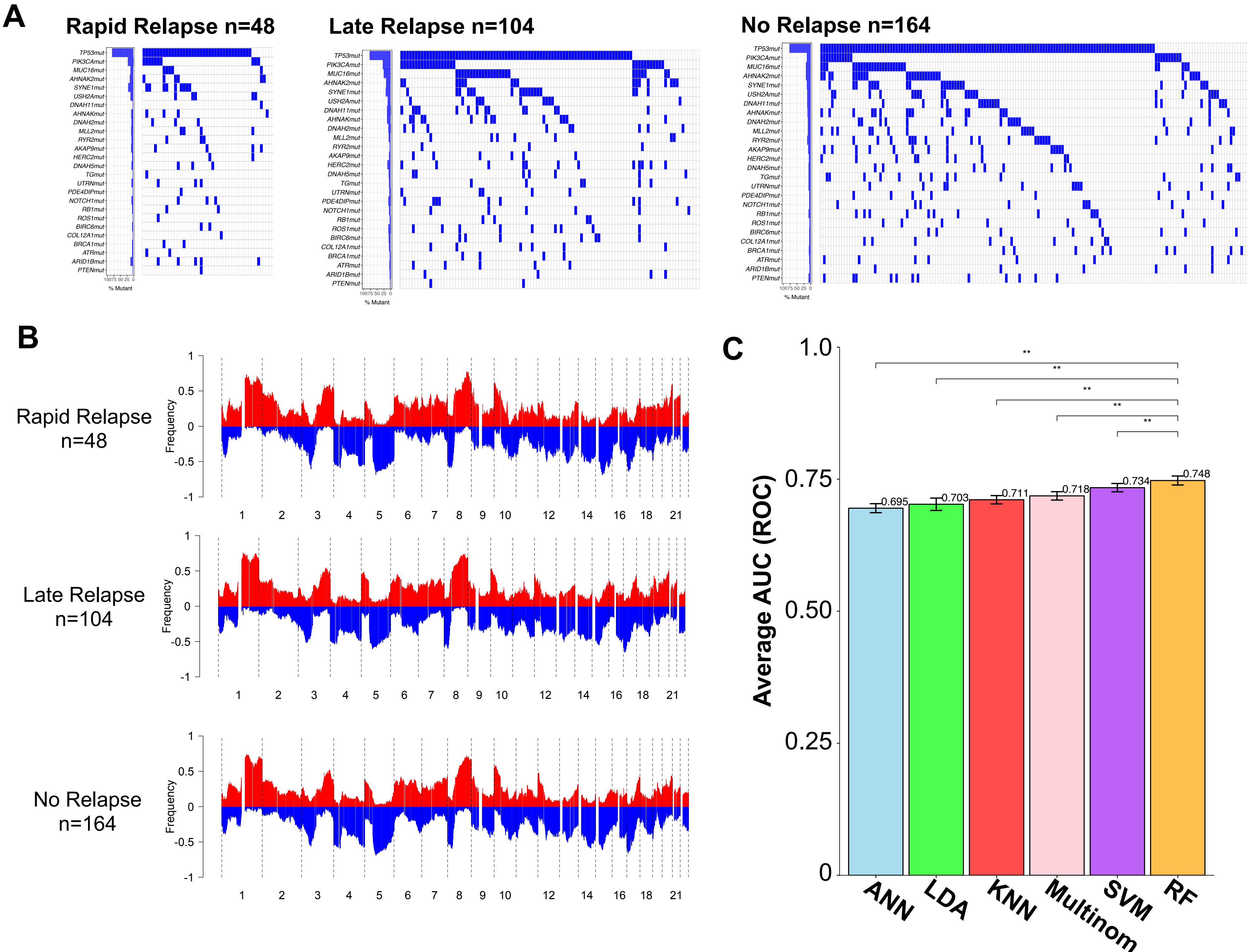
